# A zebrafish functional genomics model to investigate the role of human A20 variants *in vivo*

**DOI:** 10.1101/2020.02.23.961763

**Authors:** Daniele Cultrone, W. Nathan Zammit, Eleanor Self, Benno Postert, Jeremy ZR Han, Jacqueline Bailey, Joanna Warren, David R Croucher, Kazu Kikuchi, Ozren Bogdanovic, Tatyana Chtanova, Daniel Hesselson, Shane T. Grey

## Abstract

Germline loss-of-function variation in *TNFAIP3*, encoding A20, has been implicated in a wide variety of autoinflammatory and autoimmune conditions, with acquired somatic missense mutations linked to cancer progression. Furthermore, human sequence data reveals that the A20 locus contains ~400 non-synonymous coding variants which are largely uncharacterised. The growing number of A20 coding variants with unknown function, but potential clinical impact, poses a challenge to traditional mouse-based approaches. Here we report the development of a novel functional genomics approach that utilises the new A20-deficient zebrafish (*Danio rerio*) model to investigate the impact of *TNFAIP3* genetic variants *in vivo*. Similar to A20-deficient mice, A20-deficient zebrafish are hyper-responsive to inflammatory triggers and exhibit spontaneous early lethality. While ectopic addition of human A20 rescued A20-null zebrafish from lethality, missense mutations at two conserved A20 residues, S381A and C243Y reversed this protective effect. Ser381 represents a phosphorylation site important for enhancing A20 activity that is abrogated by its mutation to alanine, or by a C243Y mutation that associates with human autoimmune disease. These data reveal an evolutionarily conserved role for A20, but also demonstrate how a zebrafish functional genomics pipeline can be utilized to investigate the *in vivo* significance of medically relevant TNFAIP3 gene variants. This approach could be utilised to investigate genetic variation for other conserved genes.

## INTRODUCTION

In mammals, the cytoplasmic ubiquitin-editing enzyme A20, encoded by the TNFAIP3 gene, plays a key role in maintaining inflammatory homeostasis ^1^ mediated by its function to inhibit NF-κB activation ^2–6^. The medical importance of A20’s role in dampening NF-κB is highlighted by cases of germline A20 loss-of-function mutations in humans who present with severe autoinflammatory disease ^7,8^, and by the impact of A20 deletion in mice which results in spontaneous and widespread NF-κB activation, multi-organ inflammation and premature lethality ^1,9^. The emergence of A20 mutations and coding variants as causal genetic factors driving human disease ^10,11^ highlight the need to better understand A20’s functional domains controlling inflammation *in vivo*.

A20 suppresses NF-κB signalling with inhibitory activity against key molecular substrates including signalling molecules TNF receptor-associated factor 6 (TRAF6) ^9^, receptor interacting protein 1 (RIP1) ^12,13^, and the IκB kinase complex (IKK) ^14^. The A20 ovarian-tumour (OTU) domain exhibits deubiquitinating editing (DUB) activity cantered on Cys103, which cleaves activating K63-linked ubiquitin chains from RIPK1, TRAF6 and NEMO to terminate NF-κB signalling ^9,12,13,15^. The A20 zinc finger 4 (ZnF4) exhibits E3 ligase activity, adding K48-linked ubiquitin chains to RIPK1, triggering RIPK1 proteolysis ^12,16^. In addition, the C-terminal ZnF7 domain of A20 binds linear ubiquitin to non-catalytically suppress NF-κB activity ^17^. A20 is regulated at the level of gene transcription ^6^, whereby NF-κB activation induces TNFAIP3 expression ^18,19^ forming a negative feedback loop, and the inhibitory activities of A20 enzymatic sites are enhanced by IKKB-mediated serine phosphorylation at Ser381 ^20,21^.

We previously showed that coding variants that impair A20 Ser381 phosphorylation ^13^, and thereby reduce the activity of both enzymatic Cys103 and ZnF4 domains of A20 ^12,20^, exhibited a much larger *in vivo* effect in mice than mutations that disable the individual enzymatic sites Cys103 and ZnF4 ^13,21,22^. These data point towards an important role for non-catalytic Ser381 in regulating A20’s anti-inflammatory activity *in vivo*, however, the *in vivo* impact of deleting Ser381 has not been tested.

The importance of A20 in human disease, as well as the emerging functional complexities highlighted by experimental studies examining A20 mutations, coupled with the discovery of 100’s of new TNFAIP3 coding variants from human genome sequencing studies (e.g. gnomAD) highlight the need for novel approaches to investigate A20 functional domains. We hypothesised that elucidation of A20’s conserved motifs across additional species may aid the resolution of A20’s functional domains as well as highlight protein domains of relevance to human disease. Here we report the development of a novel functional genomics model that utilises the new A20-deficient zebrafish (*Danio rerio*) model to investigate the impact of TNFAIP3 genetic variants *in vivo*. Using this zebrafish paradigm we highlight the strong evolutionary conservation of A20’s role to maintain inflammatory homeostasis, and demonstrate how a zebrafish functional genomics approach can be utilized to investigate the *in vivo* significance of human TNFAIP3 gene variants. This same approach could be utilised to increase understanding of the impact of human genetic variation for other highly conserved genes.

## RESULTS

### Genomic analysis of the zebrafish *tnfaip3* locus

Many of the mammalian components of the NF-κB signalling system ^23,24^ and innate immunity ^25^ are conserved in zebrafish making this animal model a candidate *in vivo* paradigm for the analysis of A20’s key functional motifs. This concept was further supported by analysis of zebrafish genomic data which revealed gene order preservation of the TNFAIP3 locus with OLIG3 and PERP in zebrafish (Figure S1A), and a conserved and easily identifiable Topologically Associating Domain (TADs) overlapping with A20 and highly enriched in the regulatory histone mark H3K27ac (histone 3, lysine 27 acetylation) (Figure S1B), a hallmark of active promoters and enhancers ^26,27^. Furthermore, some of these conserved H3K27c marked regions overlapped with the positions of A20 SNPs identified in human genome wide association studies (GWAS) to be linked with Crohn’s disease and Systemic lupus erythematosus (Figure S1A). The *tnfaip3* zebrafish orthologue encoding a predicted 762 amino acid polypeptide comprising an N-terminal ovarian tumour (OTU) domain (identity score of 94/100), including the putative catalytic cysteine at position 103 in A20’s OTU domain ^28^, and seven zinc finger motifs in the C-terminus (identity scores of 79/100) (Figure S2). The zebrafish A20 C-terminal domain also contained zinc finger 4 (ZnF4) and ZnF7 regions, which showed multiple sequence alignment scores of 91/100 and 100/100 respectively, and are thus highly homologous to functional ZnF4 and ZnF7 domains identified in mammalian studies ^17,21^. In mammalian cells A20 expression is controlled at the level of transcription by the NF-κB pathway in response to microbial stimulation of TLR ^19^, a pathway that represents a key component of innate immunity conserved in zebrafish ^29^. Stimulation of cultured zebrafish embryos 3 days post-fertilization (dpf) with 75 µg/ml of the gram-negative bacteria product lipopolysaccharide (LPS) also resulted in the rapid induction of zebrafish (zf) zfA20 (*tnfaip3*) mRNA as well as TNF (*tnf*), zfIL-1beta (*il1b*) and zfIL-6 (*il6*) mRNA (Figure 1 A and B).

**Figure 1.**
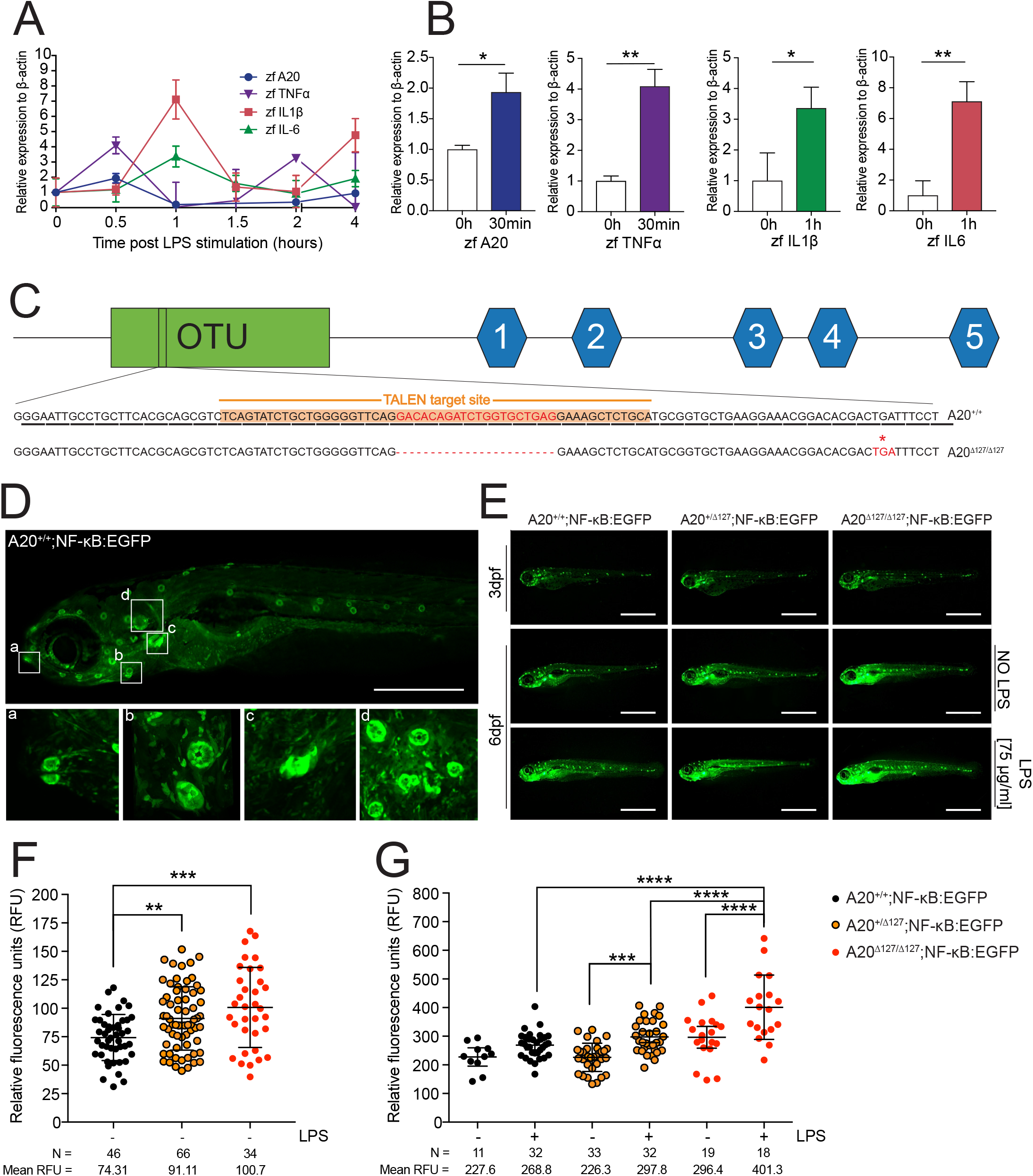
A20-deficient zebrafish fail to thrive. **A** Zebrafish were stimulated with 75 µg/ml LPS and assayed for gene expression (mRNA) at 30 min intervals. Each data point represents an N=3 pools of 10 fish each per group from three independent crosses. Data represented as mean ± SD. **B** Individual marker peak induction, and relative p-values. *P < 0.05, **P < 0.01. **C** Gene targeting of A20 highlighting the 20 bp deletion and predicted stop codon at amino acid 127, previously a Leucine. **D** Percentages of surviving A20^+/+^, A20^+/∆127^ and A20^∆127/∆127^ zebrafish at 1, 2, 3 and 8wpf, generated from N=15 A20^+/∆127^ crosses. **E** Body area measured in (pixel/inch) per zebrafish genotype at 1 and 2 wpf. Each dot represents an individual zebrafish. Data represented as mean ± SD, P-values were determined using 1way ANOVA. *P < 0.05, ****P < 0.0001. Each group passed the normality test. **F** Representative brightfield image of one A20^+/+^ (top) and A20^∆127/∆127^ (bottom) zebrafish at 2 wpf. Scale bars represent 2 mm. **G** Body to liver ratios measured in (pixel/inch) per zebrafish genotype at 1 and 2 wpf. Each dot represents an individual zebrafish. Data represented as mean ± SD, P-values were determined using 1way ANOVA. *P < 0.05, ****P < 0.0001. Each group passed the normality test. For each genotype a representative A20^+/+^, A20^+/∆127^ and A20^∆127/∆127^ 2 wpf zebrafish is shown. The scale bars represent 2 mm. **H** Immuno-fluorescence sections of livers from 2CLIP reporter fish for the A20 genotypes. The scale bars represent 300µm.

### Generation of a zebrafish model of A20 deficiency

An optimal zebrafish functional genomics model to test human A20 variants would have an A20 null genetic background. Because A20 deletion in zebrafish had not been previously reported, we utilised site directed transcription activator-like effector nucleases (TALEN) to introduce a disabling mutation within exon 2 of the zebrafish A20 locus (Figure 1C). A zebrafish line was identified that showed a 20 bp deletion in the *tnfaip3* sequence which was predicted to introduce a premature truncation at amino acid 127 (tnfaip3^gi3^, hereafter A20^∆127^; Figure 1C). Heterozygous A20^∆127^ zebrafish were bred to homozygosity and deletion of A20 was confirmed in offspring by high-resolution melting assay (HRMA), Sanger sequencing, and analysis of PCR products that differentiate A20 wild-type alleles from A20^+/∆127^, and A20^∆127/∆127^ (Figure S3). Heterozygous A20^∆127^ zebrafish were mated to generate A20^+/+^, A20^+/∆127^, and A20^∆127/∆127^ zebrafish.

### A20 deficiency is lethal in zebrafish

To validate the use of A20 deficient zebrafish for testing human A20 variants, we investigated the impact of zebrafish A20 deletion with respect to sentinel A20 phenotypes reported in mammals. The key mouse phenotypes reported in the literature include loss of organ homeostasis, gross anatomical changes (i.e. runting) and premature lethality ^1^, as well as hyper-responsiveness to TLR-induced NF-κB activity and heightened macrophage activity ^9,30^. Accordingly, A20^+/+^, A20^+/∆127^, and A20^∆127/∆127^ zebrafish were followed through time to assess survival. A20^∆127/∆127^ zebrafish presented in the expected Mendelian ratios at 1 week post-fertilization (wpf), however, by 2wpf zebrafish homozygous for A20 deletion presented with premature lethality and no A20^∆127/∆127^ zebrafish survived to three weeks post-fertilization (Figure 1D). Many A20^∆127/∆127^ zebrafish also exhibited gross abnormalities including reduced body size (Figure 1E) and evident malformations of the torso (Figure 1F). In contrast, but like mice heterozygous for A20 deletion ^1^, A20^+/∆127^ zebrafish were viable and showed no gross phenotype or lethality. Mice lacking A20 present with a rapid and early onset of systemic inflammation of the organs particularly the liver ^1^. To examine the impact of A20 deletion upon organ homeostasis A20 deficient zebrafish were crossed with 2-Color Liver Insulin acinar Pancreas (2CLIP) zebrafish, which express dsRed fluorescent protein driven by the fatty acid binding protein 10 (*fabp10*) promoter in hepatocytes ^31^. A20^∆127/∆127^:2CLIP zebrafish are phenotypically runted but also show a marked reduction in liver area disproportionate to their body size when compared to WT zebrafish (Figure 1G). Reduction in liver size was not due to a developmental defect as liver size was normal in 1 wpf A20^∆127/∆127^:2CLIP zebrafish. Confocal analysis of A20^∆127/∆127^:2CLIP zebrafish livers (Figure 1H), revealed a reduction in liver cellularity, and an increase in the frequency of caspase-3 positive hepatocytes.

### A20 limits NF-κB activation in zebrafish

To investigate NF-κB activation in A20 deficient zebrafish we crossed A20^+/∆127^ zebrafish to a NF-κB*:EGFP* transgenic zebrafish reporter line. The responsiveness of NF-κB activation in zebrafish *in vivo* is a direct result of colonisation by environmental microbiota ^24^. As shown in Figure 2A and B, on a wild type background the NF-κB*:EGFP* reporter demonstrates spontaneous activation. NF-κB driven fluorescence was found to be at its highest expression in those tissues that mostly interact with the external environment, namely the mouth, neuromasts, pharyngeal tooth and gills (Figure 2A a, b, c, d). Also, NF-κB driven fluorescence increased through time (compare 3dpf A20^+/+^ zebrafish with 6dpf; Figure 2B) most likely reflecting exposure to environmental microbiota ^24^, and this was increased further in A20^+/∆127^ and A20^∆127/∆127^ zebrafish in a gene dose manner (Figure 2B, C and D). Furthermore, the addition of LPS to zebrafish embryos further enhanced NF-κB activation and this was most pronounced in A20^∆127/∆127^ zebrafish (Figure 2C and D). NF-κB controls the expression of inflammatory response genes, and A20^∆127/∆127^ zebrafish showed increased basal expression of MPEG1 (Figure 2E), a macrophage expressed NF-κB regulated gene ^32–34^.

**Figure 2.**
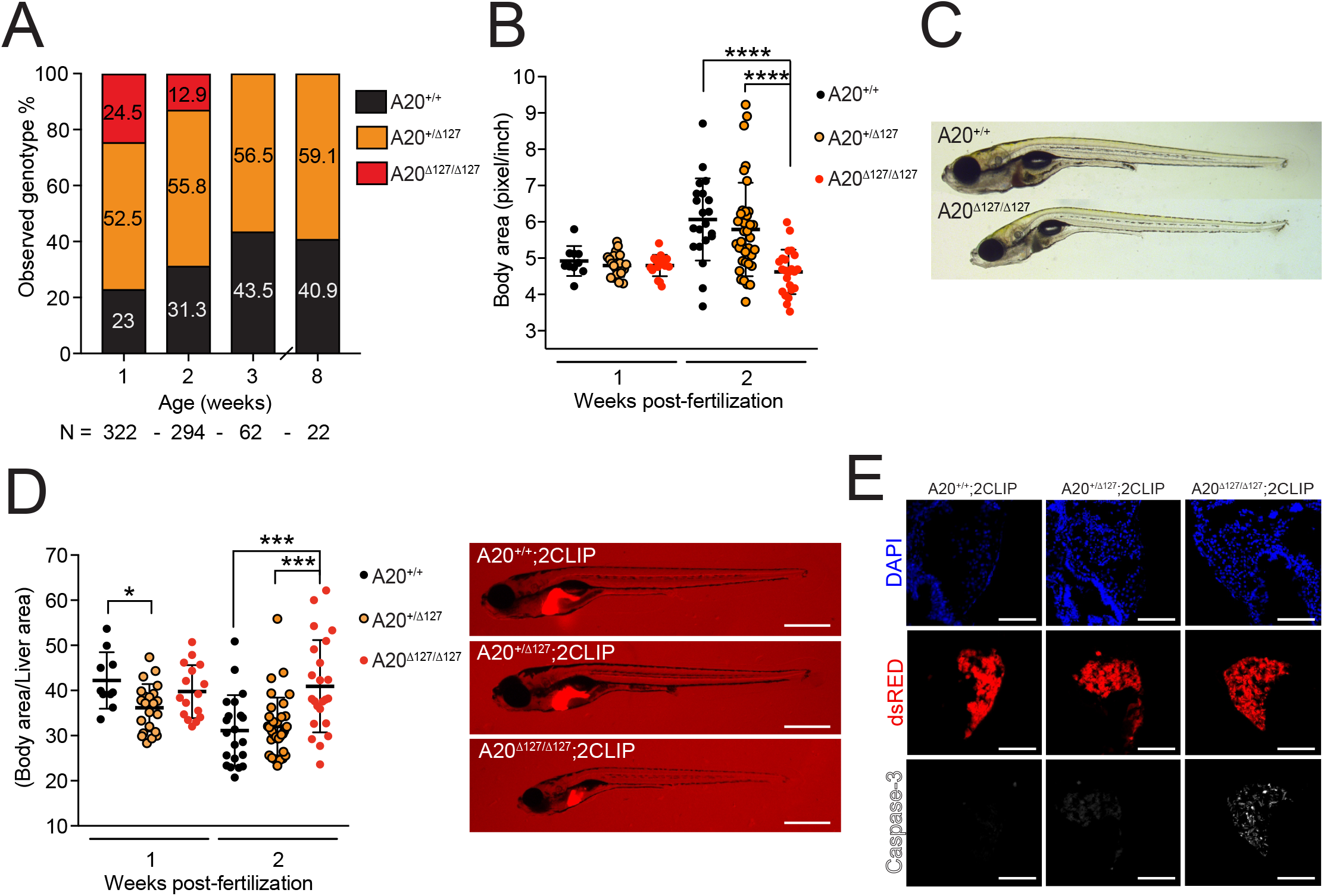
A20 is necessary to maintain inflammatory homeostasis in zebrafish. **A** Representative fluorescent image of a NF-κB:eGFP zebrafish at 6 dpf showing spontaneous NF-κB activity (fluorescence) in the (a) mouth, (b) lateral neuronmasts, (c) pharyngeal tooth, and (d) gills. Scale bar represents 2mm. **B** Representative fluorescent images of NF-κB:eGFP zebrafish of different A20 genotypes at 3 and 6 dpf ± LPS. The scale bars represent 1mm. **C-D** Cumulative data of relative fluorescence units as in (E) of NF-κB:eGFP zebrafish by A20 genotype at; (F) 3dpf without LPS stimulation and, (G), 6 dpf ± LPS. Each dot represents an individual fish. Data are representative of N=10 crosses. N per group and mean values reported below each graph. Data represented as mean ± SD, P-values were determined using 1-way ANOVA. Each group passed the normality test. **P < 0.01, ***P < 0.001, ****P < 0.0001. **E** Mpeg1 marker basal peaks at 1wpf, and relative p-values. *P < 0.05, N = 4 experiments of unstimulated zebrafish.

### Zebrafish A20-deficient macrophages are hyper responsive to inflammatory triggers

A20 loss of function mutations result in macrophage hyper responsiveness to microbial stimulation and spontaneous inflammasome activation ^9,13,17^. Tissue macrophages play a key role in sensing pathogenic microbiota in zebrafish ^33,35^. To investigate the impact of A20 deletion on macrophage homeostasis, heterozygous A20^+/∆127^ zebrafish were crossed to a macrophage reporter line whereby RFP is expressed under the control of the promotor of macrophage-expressed gene 1.1 (*mpeg1.1*) ^36^. Loss of A20 completely altered both the number and phenotype of tissue macrophages in ways consistent with increased cellular response to stimulation. The number of macrophages detected in the skin was 2-3 fold higher in A20^∆127/∆127^ zebrafish when compared to their wild type counterparts (Figure 3A and C). Furthermore, the majority of the dermal macrophages in A20^∆127/∆127^ zebrafish exhibited dendritic extensions (Figure 3A) indicative of an activated state ^37,38^. This was made further apparent by histological analysis through transverse sections of the cranium, which revealed dense infiltration of mpeg1.1 positive cells close to the dermal surface (Figure 3E). The position of the mpeg1.1 positive cells corresponded with cells at the dermal surface exhibiting bright NF-κB:EGFP reporter activity (Figure 3E).

**Figure 3.**
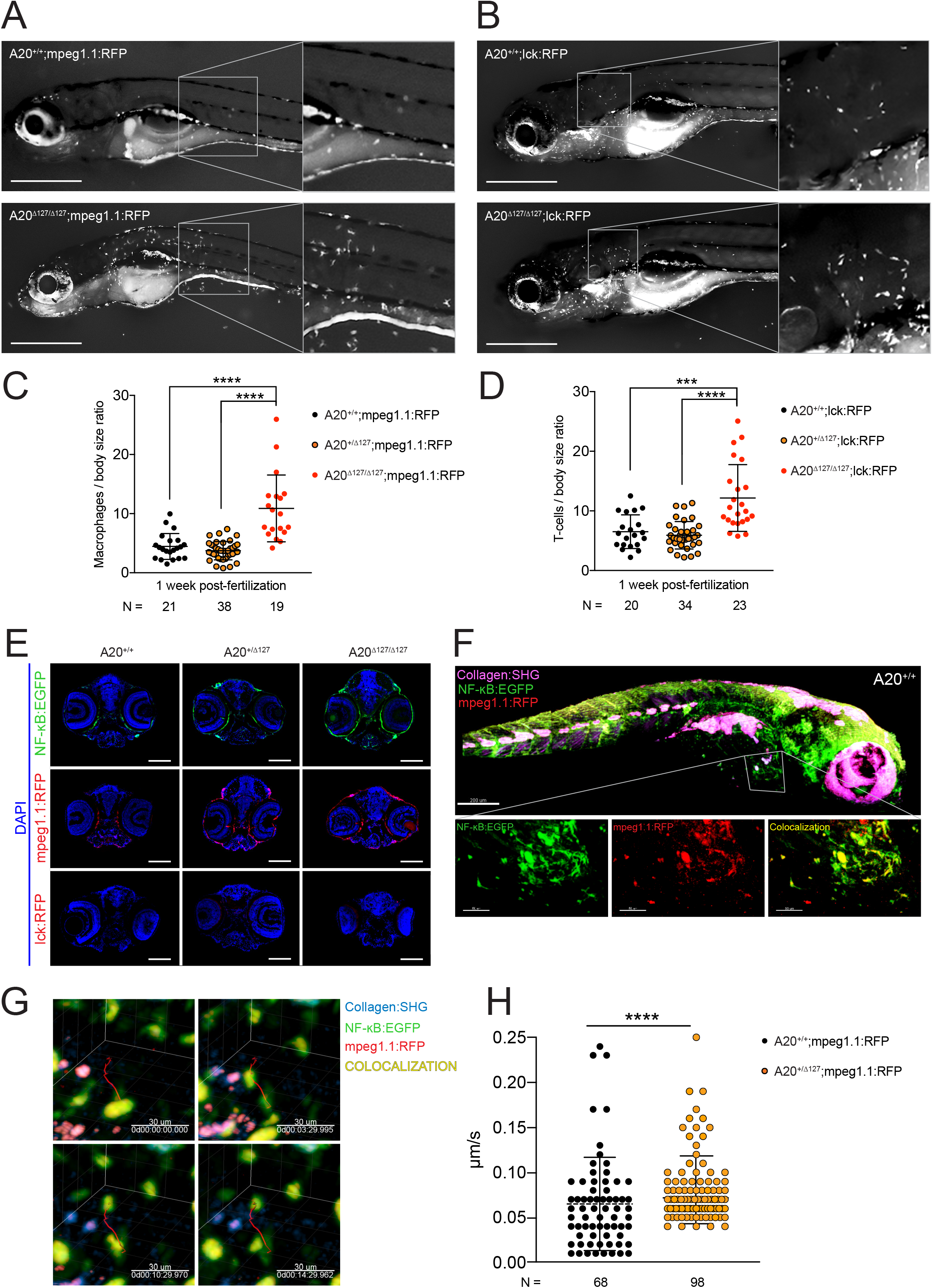
Impact of A20 deletion on macrophage activity in zebrafish. **A** Representative fluorescent image of mpeg1.1:RFP;A20^+/+^ (top) and mpeg1.1:RFP;A20^∆127/∆127^ (bottom) zebrafish at 1wpf. **B** Representative fluorescent image of lck:RFP;A20^+/+^ (top) and lck:RFP; A20^∆127/∆127^ (bottom) zebrafish at 1wpf. **C** Number of macrophages counted from images of zebrafish by A20 genotype as in (A). Data represented as mean ± SD, P-values were determined using Whelch’s t-test. **P < 0.01, ****P < 0.0001. **D** Number of T lymphocytes counted from images of zebrafish by A20 genotype as in (B). Data represented as mean ± SD, P-values were determined using Whelch’s t-test. **P < 0.01, ****P < 0.0001. **E** Representative transversal zebrafish head cryosections for a NF-κB:eGFP and mpeg1.1:RFP zebrafish for each A20 genotype at 1 wpf. The scale bar represents 75 µm. **F** Average number of macrophages at the site of injury measured every 2 hours and at 1 and 2 days post-injury. Error bands are SEM. Area under the curve was significant for A20^+/+^ x A20^+/∆127^ (****P < 0.0001) and A20^+/+^ x A20^∆127/∆127^ (***P < 0.001). **G** Macrophages from mpeg1.1:RFP positive zebrafish were imaged using *in vivo* two-photon microscopy and their track displacement, track duration and track length were quantified. Data collected from 4 A20^+/+^ and 3 A20^+/∆127^ comprising of 68 and 98 macrophages tracks respectively. Data represented as mean ± SD, P-values were determined using Mann-Whitney t-test. **P < 0.01, ****P < 0.0001. **H** Timed screenshots of a Two-Photon z-stack 15 minutes time-lapse of an mpeg1.1:RFP A20;A20^+/+^ and mpeg1.1:RFP A20;A20^+/∆127^ zebrafish. The tracks show the path taken by a representative macrophage expressing RFP. The chosen macrophages are the closest to the average tracking data.

Recruitment to sites of inflammation constitute important physiological roles for the macrophage immune compartment ^35^. Using a tail clip wound model ^39^ we found that both A20^+/∆127^ and A20^∆127/∆127^ zebrafish showed an increased recruitment of macrophages at the wound interface through time (Figure 3F). Interestingly, this analysis revealed a macrophage phenotype for heterozygous A20^+/∆127^ zebrafish post challenge whereas we did not observe a change in A20^+/∆127^ macrophage numbers in the steady state (Figure 3A and C). To investigate this further, we used two-photon imaging of single mpeg1.1:RFP reporter A20^+/∆127^ zebrafish to track individual macrophage cell movement *in vivo* in real time (Figure 3G). This analysis revealed that A20^+/∆127^ macrophages exhibited increased cell movement (Figure 3G), attributed to an increase in their average track speed, displacement and length (Figure 3H). In addition to macrophages, T cell numbers were also increased in A20 deficient zebrafish (Figure 3B and D), a phenotype comparable to A20-deficient mice which exhibit increased numbers of CD3+ lymphocytes ^1^.

### Human A20 rescues A20^∆127/∆127^ zebrafish from lethality

The phenotypic similarities between A20-deficient zebrafish and those reported for A20-deficient mice ^1,9,17^, as well as the profound inflammatory syndromes reported for human subjects with heterozygous A20 deficiency ^8^, supported utilising zebrafish as a model organism to investigate the role of A20 protein domains in regulating its function *in vivo*. For our functional analysis pipeline, we determined that rescue of A20^∆127/∆127^ zebrafish from lethality would provide the most robust readout for loss or gain of function phenotypes in various A20 genetic variants. For the functional analysis, zebrafish eggs derived from A20^+/∆127^ x A20^+/∆127^ crosses were ectopically injected with wild-type human (h)A20 expressed in the construct Ubb-hA20-P2A-EGFP that also expresses GFP. The low integration Ubb-hA20-P2A-EGFP construct was deliberately chosen to limit potential off-target effects as well as false positives ^40–42^. The zebrafish eggs were injected in a blinded fashion with the Ubb-hA20-P2A-EGFP construct and at 3wpf surviving zebrafish were assessed for their genotype and the presence of GFP fluorescence (Figure S5A). At 1 wpf A20^∆127/∆127^ zebrafish presented in the expected mendelian ratios of ~25% but exhibited a highly penetrant lethal phenotype, with no survivors at 3wpf (Figure 1D). In contrast, with ectopic expression of hA20, A20^∆127/∆127^ zebrafish comprised 9.91% of the total genotypes at 3wpf (Figure 4A). Given an expected Mendelian ratio of 25%, ectopic expression of hA20 rescued~40% of A20^∆127/∆127^ zebrafish from lethality.

**Figure 4.**
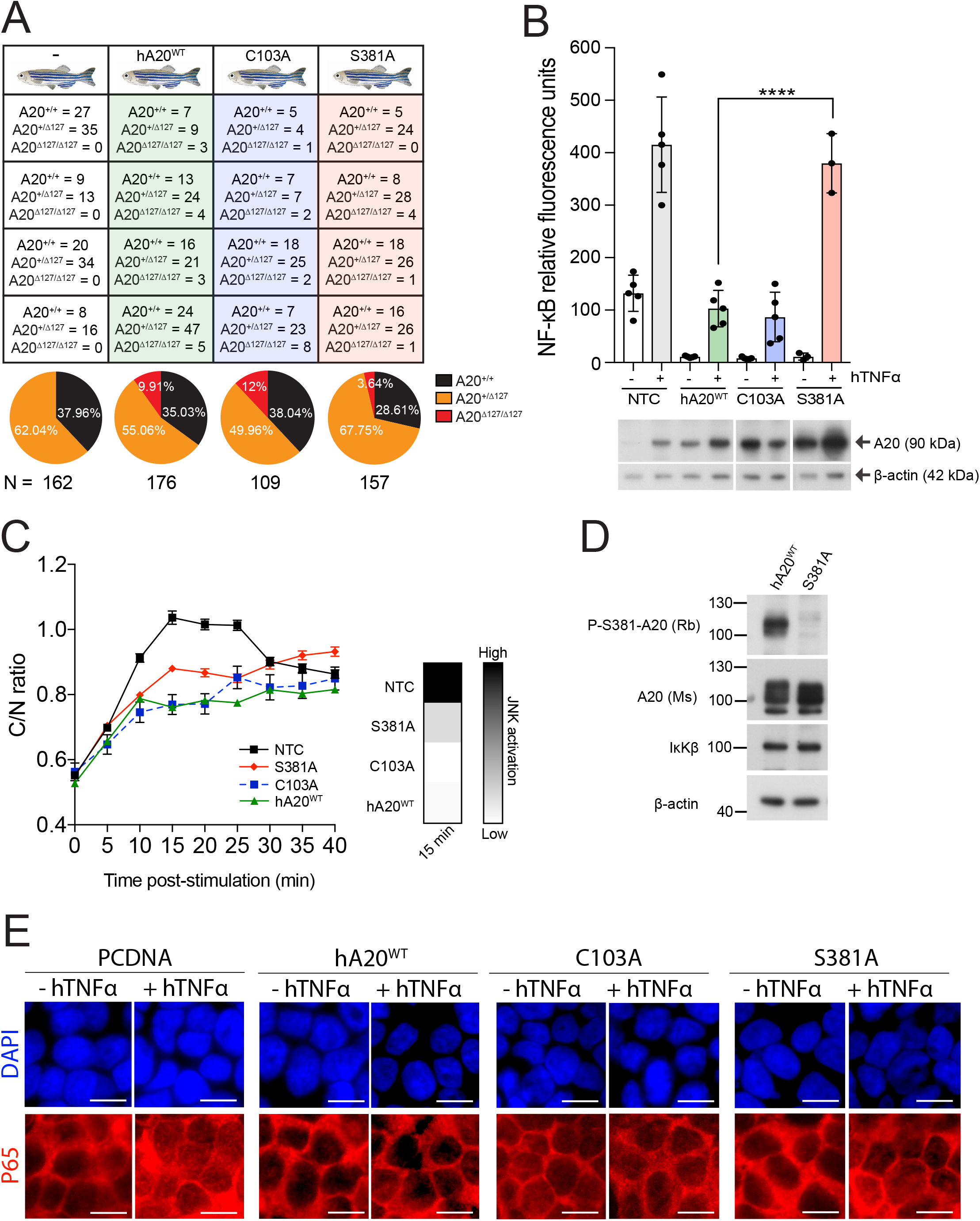
The impact of human A20 mutants on A20^∆127/∆127^ zebrafish survival. **A** Survival data for A20^+/+^, A20^+/∆127^ and A20^∆127/∆127^ zebrafish at 3 wpf with ectopic administration of WT A20 (ref-hA20) or the C103A, S381A or C243Y mutants. Numbers indicate number of genotyped survivors expressing the respective human gene variant. Each box indicates the survivors from an independent experiment. Survival frequency data is represented in the pie chart. N indicates the total number of zebrafish embryos injected for each A20 mutant. **B** NF-κB luciferase assay for HEK293 cells transiently non-transfected (NTC), or co-transfected with WT human-A20 or the C103A, S381A or C243Y mutant and treated with or without hTNFα. Data represented as mean ± SD. Each dot represents an individual experiment and are representative of N = 3-5 experiments per A20 mutant. Below is shown a representative Western blot for each A20 mutant. The displayed gel has been cropped to improve clarity and conciseness of the figure.

### Alanine substitution of Ser381 and the human mutation C243Y impair A20’s *in vivo* function

We previously showed that a series of three different A20 coding variants, all within A20’s OTU domain, showed a strong correlation between a graded loss of A20’s NF-κB inhibitory function and an increasing hyper inflammatory *in vivo* phenotype with reduced phosphorylation at Ser381^13^. The coding variant C243Y which showed the strongest *in vivo* phenotype and caused the greatest reduction in Ser381 phosphorylation was originally identified in a four generation Japanese family with hereditary autoinflammatory disease ^7^. *In vitro* biochemical studies show that IκB kinase beta (IKKβ)-dependent phosphorylation at Ser381 is required for A20’s optimal NF-κB and JNK suppressive function ^13,20,43^ (Figure 4B; Figure S5B and C). Both Ser381 and Cys243 are conserved in zebrafish (Figure S2) allowing us to test the impact of alanine substitution at Ser381, and the impact of the disease causing variant C243Y on the ability of human A20 to rescue A20^∆127/∆127^ zebrafish from lethality.

As shown in Figure 4B, A20^∆127/∆127^ embryos that received either S381A or C243Y expressing constructs were found in reduced numbers when compared to A20^∆127/∆127^ embryos that received wild type hA20-expressing constructs. Specificity for the impact of S381A and C243Y on A20’s *in vivo* function was shown by ectopic injection of another OTU domain mutant, namely the C103A DUB mutant ^13,21,22^, which exhibited no negative impact on hA20’s ability to block NF-κB or JNK signalling *in vitro* (respectively Figure 4B; Figure S5B and C), nor A20’s ability to rescue A20^∆127/∆127^ embryos from lethality (Figure 4A). The *in vivo* rescue properties of individual A20 mutants did not relate to differences in protein stability, as shown by Western blot analysis of A20 protein levels when expressed in cell lines (Figure 4B) ^13^, nor differences in the levels of transgene expression *in vivo* as measured by the levels of GFP fluorescence which is co-expressed with A20 in the Ubb-hA20-P2A-EGFP construct (Figure S5A).

## DISCUSSION

A20 inhibits NF-κB activation in mammals in response to TLR signals ^1,9^, and uncontrolled NF-κB activation results in excessive inflammation and lethality ^44^, yet there is little information regarding the *in vivo* functional role of A20 beyond studies in mammalian models. An A20-like protein sequence has been identified in the purple sea urchin and amphioxus ^45^ and in the lamprey genomes ^46^ and here we identify a functional A20 homologue in zebrafish. In contrast, we did not find a homologous A20 sequence in the genomes of lower invertebrates including *Drosophila*. Similar to mammalian cells, and as shown here for zebrafish, A20 expression in amphioxus is also regulated by LPS ^19,45^ and A20 modulates NF-κB signalling utilizing a ubiquitin editing function ^13,21,45^. However, the *in vivo* functional significance of A20 in amphioxus has not been tested ^45^. Reminiscent of mouse data ^1,9^, we found that A20 deficiency in zebrafish was lethal and remarkably re-introduction of human A20 could rescue a proportion of A20-deficient zebrafish from lethality. We interpret these data to suggest that A20 arose during chordate evolution to regulate inflammatory homeostasis and prevent lethality, most likely in response to environmental microbial stimuli. As the major components of NF-κB signalling in mammals are conserved to *Drosophila* ^13,47^ it is unclear as to why A20-dependent control of NF-κB signalling was required for chordate evolution. Also, in mammalian cells A20 has been shown to regulate diverse signalling cascades that control inflammation and cell death in both hematopoietic and non-hematopoietic cell lineages ^10,48^. Further investigation of A20’s function in non-mammalian systems like zebrafish will reveal to what extent these critical mammalian functions extend to other chordates and aid our understanding of the evolutionary significance of A20.

Gene targeting in mice to either introduce an alanine at Cys103 (C103A) that eliminates biochemical OTU DUB activity ^21^, or a mutation that destroys the ZnF4 E3 ligase activity of A20 ^22^, resulted in surprisingly little or no impact on lipopolysaccharide (LPS) responses and inflammatory homeostasis *in vivo* ^22,49^, contrasting the lethal hyper-inflammatory phenotype of A20-deficient mice ^1,9^. These findings indicate compensatory activity of the Cys103 and ZnF4 catalytic domains towards each other, and redundancy in the mechanism by which the OTU domain regulates NF-κB activation which may include both catalytic and non-catalytic roles ^12,14^. Previous biochemical studies have shown that A20’s NF-κB inhibitory activity requires phosphorylation at Ser381 by IκB-kinase ^20,21^. We previously provided indirect evidence for the importance of Ser381 *in vivo* ^13^. We have recently demonstrated how a series of A20 coding variants that cause a decrease in phosphorylation at Ser381 also decrease A20’s NF-κB inhibitory activity in a graded manner, with a corresponding loss in the ability of A20 to control inflammatory homeostasis *in vivo*. Here, using a new zebrafish functional genomics approach, we show evidence that Ser381 phosphorylation is needed for A20’s optimal anti-inflammatory function *in vivo*, as targeted mutation of Ser381 impairs the ability of A20 to rescue A20^∆127/∆127^ zebrafish from lethality. Prevention of Ser381 phosphorylation may disable both OTU DUB and ZnF4 ubiquitin editing functions most likely explaining to the highly penetrant phenotype seen for both zebrafish and mice ^13^ compared to the individual inactivation of catalytic domains ^22,49^. Further studies are required to understand how phosphorylation at Ser381 impacts A20’s function but one possibility is that phosphorylation causes changes to A20 structure facilitating interactions between A20’s distinct ubiquitin binding and ubiquitin editing domains.

Mutations in the A20 ZnF7 non catalytic NF-κB inhibitory domain cause spontaneous macrophage inflammasome activation and arthritis in mice ^17^. The cell type specific and different phenotype of ZnF7 mutant mice to that reported for mice with reduced Ser381 phosphorylation ^13^ suggests that A20 regulatory functions can be distinguished from one another. Ser381 may regulate both OTU and the ZnF4 dependent-ubiquitin editing functions, whilst ZnF7 regulates linear ubiquitin chains outside the aforementioned mentioned domains. This highlights how specific A20 functional domains are important within particular physiological and/or cellular contexts. Understanding how specific molecular changes in A20 function link to cellular phenotypes may provide a pathway to understand the seemingly broad association of A20 to many different tissue specific autoimmune diseases identified through GWAS ^10^ and the broad clinical phenotypes emerging in patients identified with A20 haploinsufficiency ^43,50,51^.

A20 deletion results in severe autoinflammatory disease in humans ^30,52–55^. The severe impact of mutating Ser381 on A20 function shown here, and the discovery of an ever increasing number of new A20 coding variants through human sequencing studies ^13^, raise the question as to the existence of further human coding variants which may present with highly penetrant phenotypes and contribute to disease. The conservation of A20 in zebrafish allows us to construct a functional genomics pipeline to investigate the impact of medically and scientifically important A20 coding variants *in vivo*. The discovery ^7^ of the cysteine to tyrosine substitution at position 243 (C243Y) in a four generation Japanese family with a Behcet’s like disease, together with the high level of sequence homology across the mammalian and zebrafish OTU domain which included this residue, provided an opportunity to test the sensitivity of the zebrafish pipeline. In this zebrafish model the C243Y mutation caused a reduction in A20’s ability to rescue homozygous A20-null zebrafish from lethality. This *in vivo* proof of concept data in the zebrafish model showing that C243Y is a reduction of function variant is supported by the human genetic data ^7^, as well as biochemical and mouse data ^13^. A20 variants like C243Y that modify phosphorylation at Ser381 therefore represent a novel class of variants with medically relevant disease-causing potential.

We highlight the potential of the zebrafish system to advance our knowledge of medically significant A20 gene variants by taking advantage of regions of high homology. In addition to shedding light on the functional significance of A20 coding variants, the conservation of the zebrafish A20 genomic locus may also aid the elucidation of non-coding variants. GWAS studies associate A20 with multiple autoimmune conditions and many identified SNPs are non coding ^10^. Analysis of conserved trans regulatory regions in the A20 locus marked by active methylation marks revealed an overlap with two SNPs associated with Crohn’s disease (rs7753394; rs7773904) (GRCh38) and one associated with Systemic lupus erythematosus (rs10499197) ^56–58^. As these SNP’s lay in putative enhancer regions they may alter A20 expression levels thereby contributing to disease. The development of a novel functional genomics pipeline that utilises the new A20-deficient zebrafish model provides a new tool to investigate the impact of TNFAIP3 genetic variants *in vivo*. Understanding the underlying genetic code of A20 will have relevance for understanding human disease mechanisms and the development of targeted drug therapies. This same approach could be utilised to increase understanding of the impact of human genetic variation for other highly conserved genes.

## ACKNOWLEDGMENTS

We thank the Biological Testing Facility at the Garvan Institute of Medical Research for fish care, and David Zahra (Genomic Engineering, Garvan Institute of Medical Research) for helpful advice with design of TALEN constructs, cloning and site directed mutagenesis of A20 mutants. We thank Dr John Rawls (Department of Cell and Molecular Physiology, University of North Carolina, Chapel Hill) for the NF-κB*:EGFP* transgenic construct. D.C. and N.W.Z were each supported by an Australian Postgraduate Award and N.W.Z. is an International Pancreas and Islet Transplant Association (IPITA) Derek Gray Fellow. The research was supported by National Health and Medical Research (NHMRC) grants to S.T.G. (1130222, 1189235) and K.K. (1130247); as well as grants to S.T.G. from the NIH (DK076169) and the Australian Juvenile Diabetes Research Foundation (3-SRA-2018-604-M-B). S.T.G. is NHMRC Senior Research Fellow (1140691).

## AUTHOR CONTRIBUTIONS

D.C., B.P., D.H. and S.T.G generated the Tg(NF-κB, c-FOS:EGFP)^GI1^ and D.C. and S.T.G. generated the tnfaip3^GI2^ zebrafish lines. Genomic analysis of the zebrafish *tnfaip3* locus was conducted by O.B.. All *in vivo* zebrafish studies were conducted by D.C. Cloning of human A20 and A20 mutants, Western blotting experiments and NF-κB reporter assays were conducted by D.C. and N.Z.W. JNK reporter assays were conducted by D.C., J.Z.M.H. and D.R.C.. J.B. and T.C. carried out intravital imaging. Immunohistochemistry was conducted by J.W., D.C and E.S.. Zebrafish expertise and essential experimental advice for use of zebrafish reporter lines was provided by K.K. D.C., N.W.Z., B.P., J.Z.R.H., J.W., D.C., K.H., T.C., D.H. and S.T.G. critically assessed data. D.C. and S.T.G. co-wrote the manuscript. S.T.G. devised and led the experimental plan and is guarantor of the study.

## MATERIALS AND METHODS

### Zebrafish

Fertilized eggs were collected from zebrafish (*Danio rerio*) group matings and incubated at 28.5 °C at a density <60 embryos in ~25 mL E3 egg water. Adult zebrafish were maintained under standard conditions at 28.5 °C and pH 7-8 at a density of 5-10 fish L^−1^ under a photoperiodic cycle of 14/10 light/dark hours. Genotyping was performed by fin clipping and HotSHOT DNA extraction ^59,60^ followed by High Resolution Melting assay. Animal studies were approved by the Garvan/St Vincent’s Animal Ethics Committee. All procedures performed complied with the Australian Code of Practice for Care and Use of Animals for Scientific Purposes.

### Genomic structure of A20

HiC data (40Kb resolution) corresponding to H1 cells were visualized in the 3D genome browser ^61^ (hg19 genome assembly). Layered H3K27ac ChIP-seq data from seven cell lines (GM12878, H1-hESC, HSMM, HUVEC, K562, NHEK, NHLF), available through the UCSC genome browser ^62^ were generated by the ENCODE consortium (2012). Zebrafish 24hpf embryo H3K27ac ChIP-seq reads were mapped by Bowtie ^63^ to unique genomic positions, followed by duplicate removal. Mapped ChIP-seq reads were visualized in WIG format as previously described ^64^ utilising the UCSC genome browser.

### *Tnfaip3* knockout line

A pair of TALENs targeting zebrafish *tnfaip3* (A20) exon 2 were generated using “PLATINUM Gate TALEN Kit” (Addgene, #1000000043). 0.2 ng of mRNA encoding the TALEN pair was delivered into the cytoplasm of wild-type zebrafish embryos at the one-cell stage. Founders were genotyped using the following primers: zfA20_Fw: AGTATCTGCTGGGGGTTCAGGA and zfA20_Rev: CACCGCATGCAGAGCTT. Subsequently a founder line was identified harbouring a 20 bp deletion in exon 2 with a predicted stop codon at amino acid 127. The *tnfaip3*^dGI4^ A20-deficient line was maintained in the heterozygous state (tnfaip3^∆127^/tnfaip3^+^) and crossed to the zebrafish reporter lines described below.

### Zebrafish reporter lines

The lck:RFP (TgBAC(*lck*:RFP)^vcc4^) and mpeg1.1:RFP (TgBAC(*mpeg1.1*:RFP)^vcc7^) reporter lines have been described previously ^36^. The lymphocyte protein tyrosine kinase (*lck*) promoter targets gene expression to T cells whereas macrophage expressed gene 1.1 (*mpeg1.1*) promoter targets gene expression to macrophages.

The Tg(ins:dsRed)^m1081^;Tg(fabp10:dsRed;ela3l:GFP)^gz12^ line, known as 2-Color Liver Insulin acinar Pancreas (2CLIP) fish line, has been described previously ^65^. This fish express dsRed fluorescent protein both in the hepatocytes, driven by the fatty acid binding protein 10 (*fabp10* gene), and in the islets of the endocrine pancreas, driven by the insulin promoter (*insulin* gene). The exocrine pancreas instead expresses GFP driven by the pancreas specific transcriptor factor 1a (*ptf1a* gene). The Tg(*NF-κB, c-FOS*:EGFP)^GI5^ zebrafish line was generated using the Tol2 system to insert a DNA fragment containing six repeats of an NF-κB binding site in tandem with a minimal c-Fos promoter that drive the expression of eGFP, followed by an SV40 polyadenylation signal sequence (SV40pA) ^24^.

### Fluorescence quantification through ImageJ

Live zebrafish were placed in a gel solution (1x E3 solution, 3% methyl cellulose, 5% Tricaine), in a 150 mm petri dish lid. Fish were placed on the left side up. Images were processed using ImageJ (v.1.50b). In the case of the NF-κB:eGFP line fluorescence measurements, an outline of the fish was drawn by hand, excluding the yolk sack due to its auto fluorescence. Measurement values of area (pixel/inch) and integrated density (area × mean grey value) were recorded for each of the measured fish. Background value was subtracted to the fish intensity value.

### NF-κB and JNK reporter assays

The NF-κB reporter assay was conducted exactly as described previously ^19^. In brief, human Embryonic Kidney 293T (HEK293T) cells ^66^ were cultured in Dulbecco’s Modified Eagle’s medium (DMEM) and incubated at 37°C in 5% CO_2_ and kept at a passage number between 21-40 for experiments. Cells were transfected using Lipofectamine 2000 (ThermoScientific #11668019) with β-galactosidase (β-gal) and NF-κB.Luc with either ‘empty’ pcDNA3.1 vector (NTC; non template control) or pcDNA3.1 encoding wild-type human-A20 or either the A20 mutant Cys103Ala or Ser38Ala. HEK293T cells were stimulated with 300 U/ml of recombinant human TNFα (RandD Systems, Minneapolis, MN; #210-TA) for 8 hours. To assess the effect of A20 mutants on JNK signalling, HEK293T cells which constitutively expresses mCherry Kinase Translocation Reporter (mCherryKTR) which translocates from the nucleus to the cytoplasm when phosphorylated by JNK were used (de la Cova, Townley et al. 2017). JNK activity was determined by measuring the ratio between nucleus and cytoplasmic mCherry in cells transiently transfected with human-A20 mutants (using JetPRIME reagent) following human TNFα stimulation. Each experiment included automated cell imaging (Thermo Scientific Cellomics Arrayscan machine) of ~150, 000 to ~350, 000 individual cells per well in duplicate wells over a period of 60 minutes (5 minutes intervals).

### Generation of A20 mutants for ectopic expression in zebrafish

Two A20 mutants were generated for this study. Mutations were introduced into cDNA encoding full length human A20 by site directed mutagenesis to cause an alanine substitution at position Cys103 (C103A) and at Ser381 (S381A). Wild type A20 or either mutant C103A or S381A A20-cDNA were cloned into the expression cassette Ubb-[A20 insert]-P2A-EGFP and ~100 ng/µl of DNA was co-injected with I-SceI meganuclease to promote random integration in the zebrafish genome. Glass needles for zebrafish eggs injections were prepared from glass capillary tubes with filament (ø1 mm × 90 mm) (WPI #1B100F-3) using a NARISHIGE dual-stage micropipette puller PC-10 machine. The injections were all performed between the one and two-cell stage of development.

### Two-photon intravital microscopy

Two-photon imaging was performed using an upright Zeiss 7MP two-photon microscope (Carl Zeiss) with a W Plan-Apochromat 20×/1.0 DIC (UV) Vis-IR water immersion objective. Four external NDDs were used to detect blue (SP 485), green (BP 500-550), red (BP 565-610) and far red (BP 640-710). High repetition rate femtosecond pulsed excitation was provided by a Chameleon Vision II Ti:Sa laser (Coherent Scientific) with 690-1064nm tuning range. We acquired 3µm z-steps at 512×512 pixels and resolution 0.83µm/pixel at a frame rate of 10 fps and dwell time of 1.27 µs/pixel using bidirectional scanning. Intravital two-photon microscopy was performed by embedding 1wpf zebrafish in 1% low melting point agarose in E3 water and 0.4% Tricaine to limit the overall tissue drifting and motility of the fish.

### Image processing and data analysis

Raw image files were processed using Imaris (Bitplane) software. A Gaussian filter was applied to reduce background noise. Tracking was performed using Imaris spot detection function to locate the centroid of cells. Motility parameters such as cell displacement (or track length calculated as the total length of displacements within the track) were obtained using Imaris Statistics function.

### Statistical analysis

Data were analysed as appropriate using a nonparametric Mann-Whitney test, a Whelch t-test, Area under the curve (AUC) or a one-way ANOVA test. Statistical analyses were carried out using GraphPad Prism v7.0. In all cases, the significance threshold was set at *p* ≤ 0.05.

## ONLINE SUPPLEMENTAL MATERIAL

**Figure S1.**
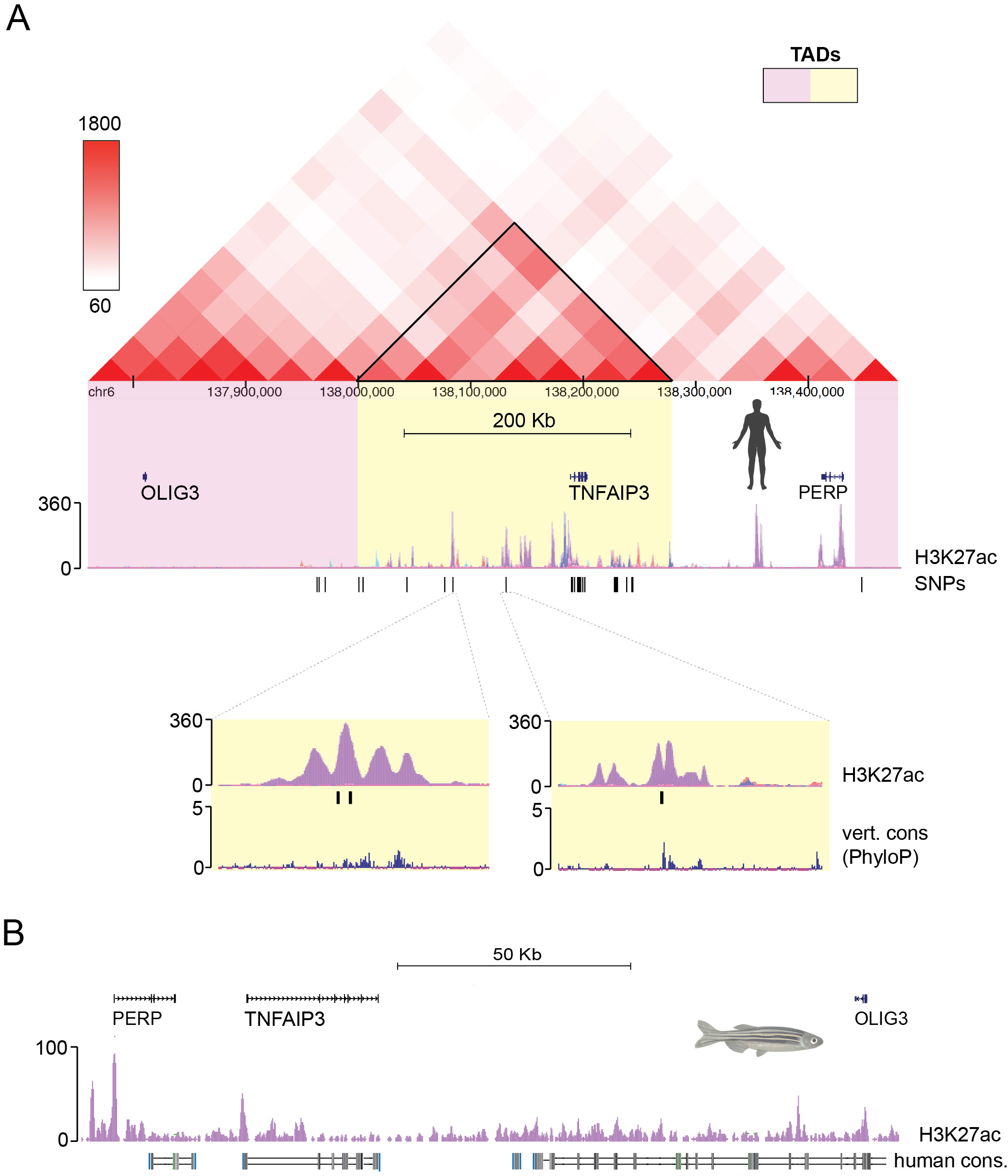
Genomic structure of the TNFAIP3 (A20) locus. **A** HiC data (H1 cells) visualized over the A20 locus, overlapping A20 SNPs and H3K27ac ChIP-seq signal. Insets correspond to three evolutionarily-conserved (PhyloP, vertebrate conservation) A20 SNPs, which coincide with the regulatory H3K27ac mark. The alternating yellow and pink shaded areas are predicted Topologically associated domains (TADs), whereas the heatmap (white-red) represents normalised interaction frequency, as described in (Wang, Song et al. 2018). The SNP IDs for SNPs with insets in the figure are (from left to right on the figure): **a)** chr6:138085249-138085249 (rs7753394), **b)** chr6:138085366-138085366 (rs7773904), **c)** chr6:138132517-138132517 (rs10499197). **B** H3K27ac ChIP-seq (24 hpf embryos) visualized over the A20 locus. Dark boxes represent human-fish DNA sequence conservation.

**Figure S2.**
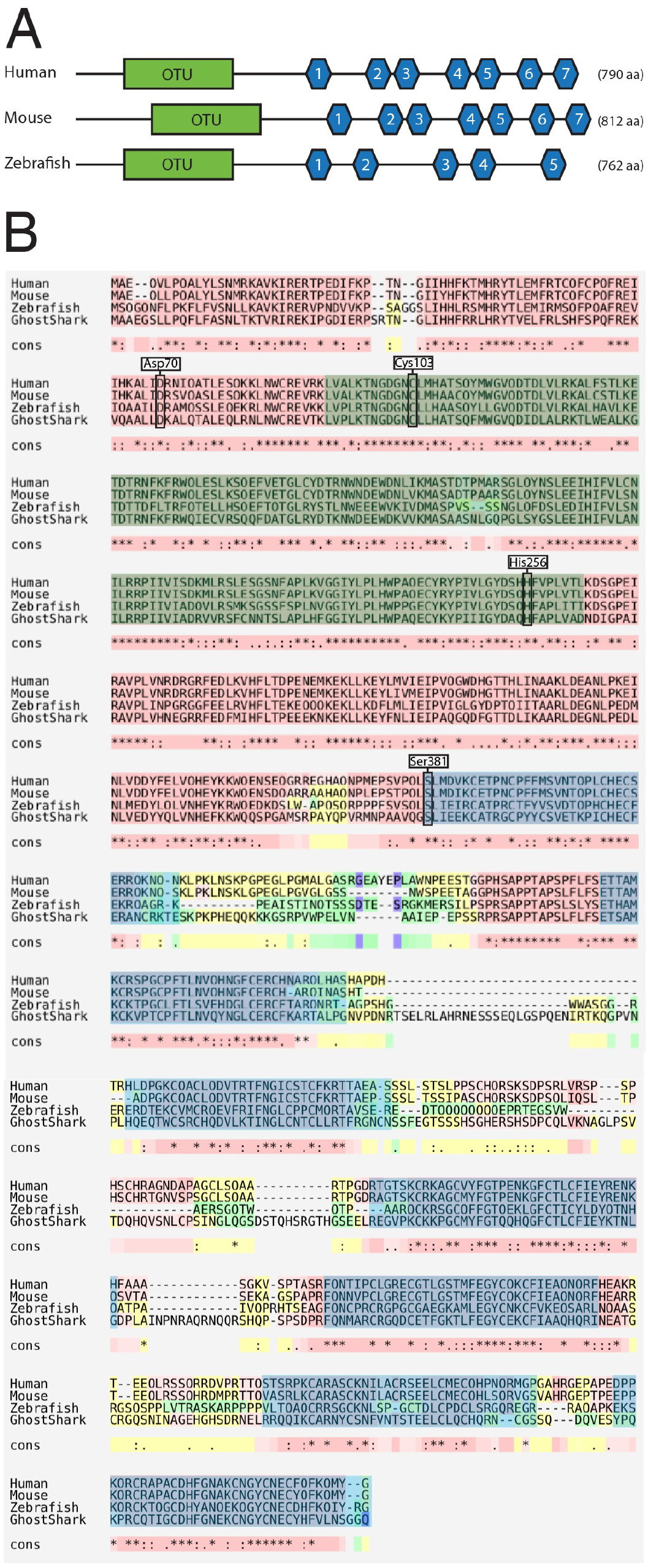
Protein homology of TNFAIP3 across multiple vertebrate lineages. The consensus sequence (cons) below each amino acid reports an “*” (asterisk) when all the amino acids are the same, a “:” (colon) indicates conservation between groups of strongly similar properties and a “.” (period) indicates conservation between groups of weakly similar properties. The amino acids comprising the OTU domain catalytic triad (Asp70, Cys103, His256) and position Ser381 are also shown. Note these two protein motifs are explicitly conserved across species.

**Figure S3.**
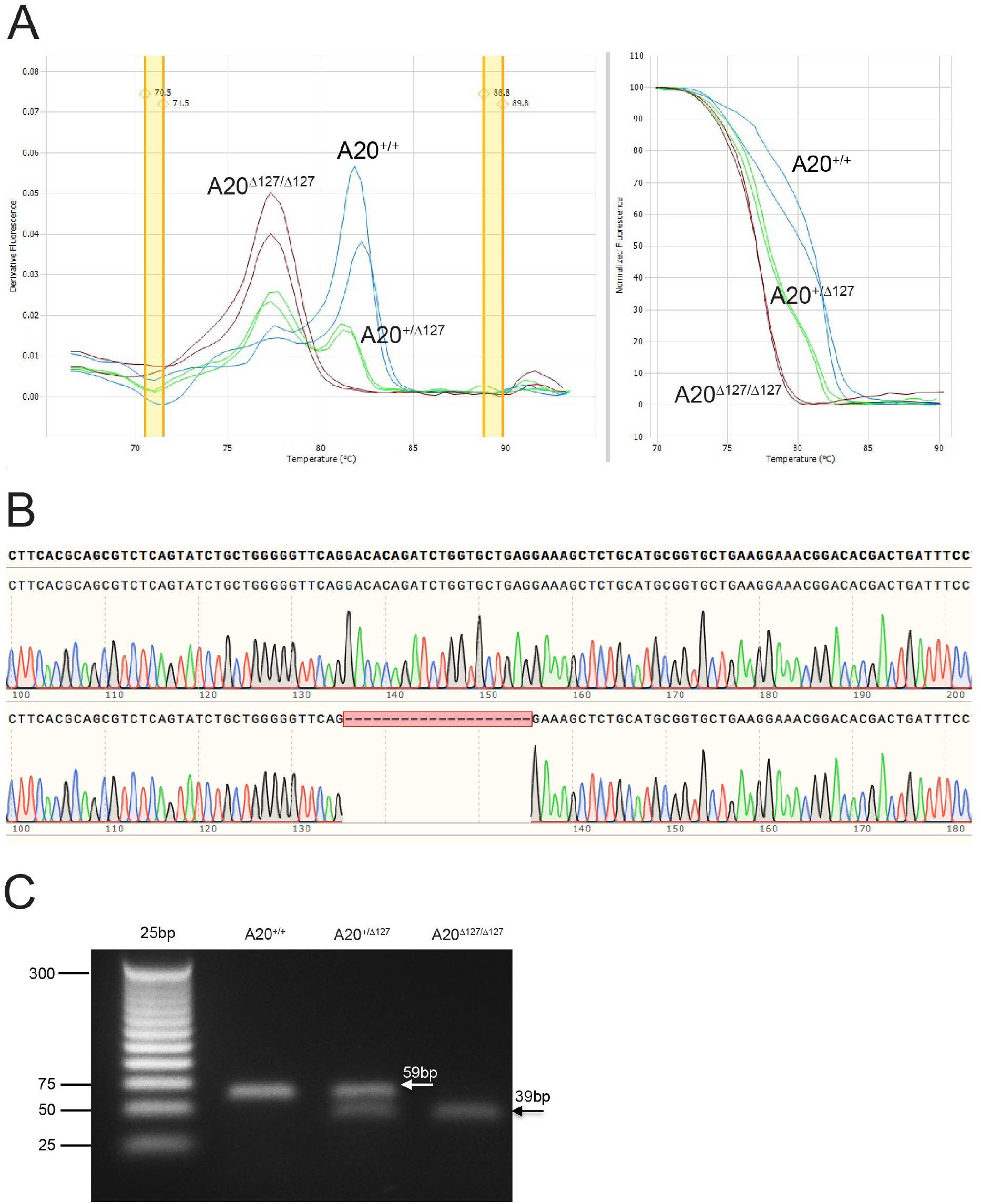
Validation of A20 deletion in TALEN targeted zebrafish. **A** HRMA curves (derivative on the left and raw-melt on the right) showing the three possible genotypes arising from a cross between two A20^+/∆127^. The trace for a zebrafish carrying either A20 wild type alleles (Blue), or heterozygous (Green) or homozygous (Red) for the mutant A20 allele are shown. **B** Shows sequence trace and alignment for a zebrafish carrying either the A20 wild type A20 allele or the deletion. **C** Shows PCR amplification of the TALEN targeted region of the A20 gene for A20^+/+^, A20^+/∆127^ and A20^∆127/∆127^ zebrafish. The displayed gel has been cropped to improve clarity and conciseness of the figure.

**Figure S4.**
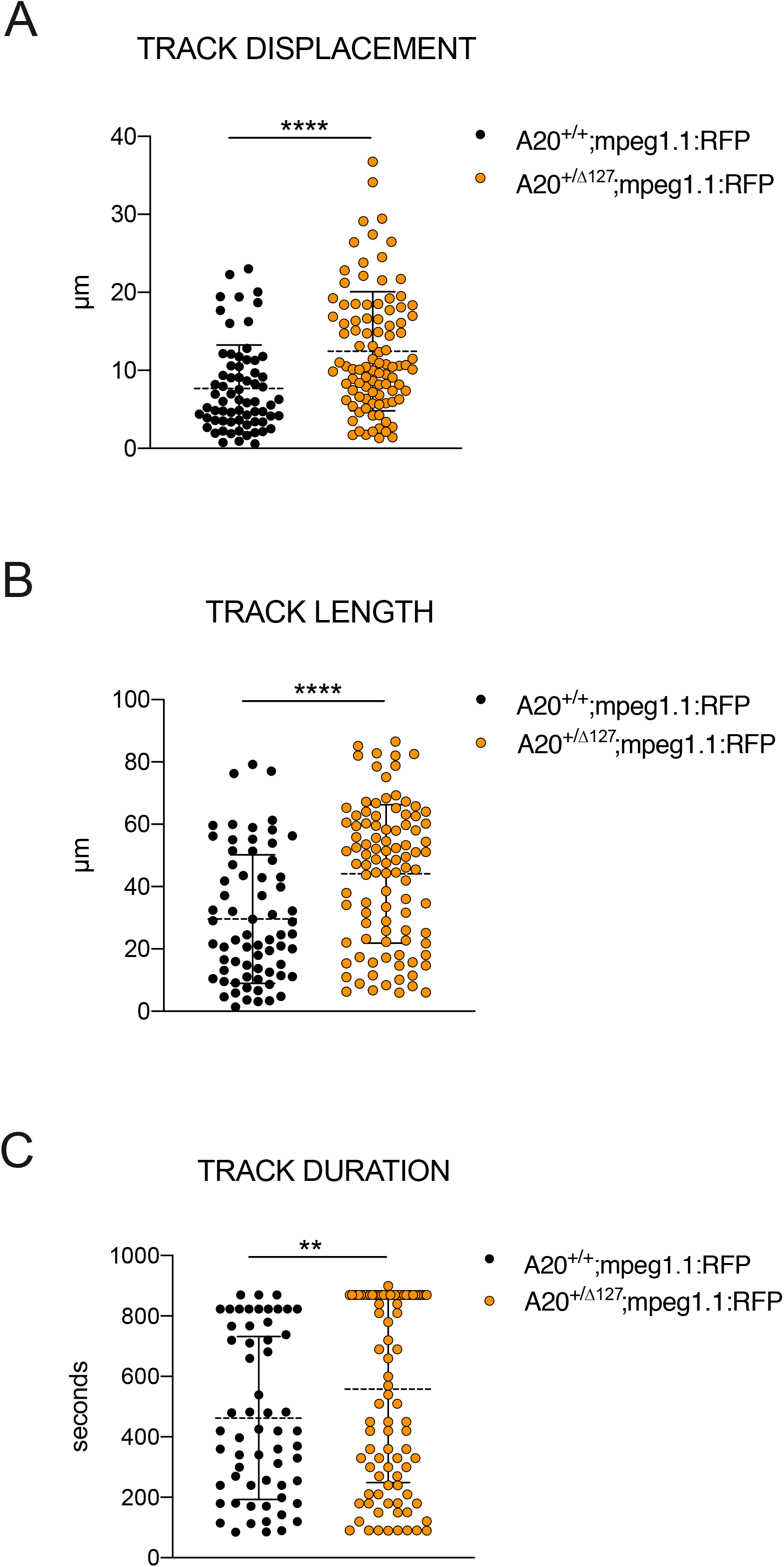
Co-localisation of NF-κB and MPEG1 reporter signals in zebrafish. Two-photon microscopy 3D-volume of a double reporter (NF-κB:eGFP;mpeg1.1:RFP) A20^+/+^. Below are shown each individual channel and the merge to demonstrate colocalization of the two markers and highlighted by dotted white lines circling the cells where colocalization occurrs. Image supplemented by video file (Video S3).

**Figure S5. Control Data for A20-dependent JNK inhibition and *in vivo* gene expression**

**A** Relative fluorescence units, corrected for fish size. No statistical significance was observed between groups of fish injected with different human A20-EGFP constructs. **B** Impact of A20 variants in mammalian cells. JNK activation kinetics for HEK293 cells transiently non-transfected (NTC), or co-transfected with WT human-A20 or the C103A, S381A or C243Y mutant and treated with or without hTNFα for 1 hour and quantified at 5 minutes intervals. Data shows the ratio of Cytoplasm over Nuclear (C/N) JNK activation, represented as mean ± standard error. **C** Data from the 15 min time point for all A20 variants is represented in heat map format. Statistical analysis carried as Area Under the Curve (AUC) compared to WT hA20. WT and NTC = ****P < 0.0001, WT and S381A or C243Y = *P < 0.05. The data on this graph have been generated by three independent experiments.

